# Histone H3 dopaminylation in ventral tegmental area underlies heroin-induced maladaptive plasticity

**DOI:** 10.1101/2021.03.26.437235

**Authors:** Sasha L. Fulton, Swarup Mitra, Ashley E. Lepack, Jennifer A. Martin, David M. Dietz, Ian Maze

## Abstract

Persistent transcriptional eventsamples were incubated on ice for 30 min and then centrifugeds in ventral tegmental area (VTA) and other reward relevant brain regions contribute to enduring behavioral adaptations that characterize substance use disorder (SUD). Recent data from our laboratory indicate that aberrant accumulation of the newly discovered histone post-translational modification (PTM), H3 dopaminylation at glutamine 5 (H3Q5dop), contributes significantly to cocaine-seeking behavior following prolonged periods of abstinence. It remained unclear, however, whether this modification is important for relapse vulnerability in the context of other drugs of abuse, such as opioids. Here, we showed that H3Q5dop plays a critical role in heroin-mediated transcriptional plasticity in midbrain. In rats undergoing abstinence from heroin self-administration (SA), we found acute and persistent accumulation of H3Q5dop in VTA. By attenuating H3Q5dop during abstinence, we both altered gene expression programs associated with heroin withdrawal and reduced heroin-primed reinstatement behavior. These findings thus establish an essential role for H3Q5dop, and its downstream transcriptional consequences, in opioid-induced plasticity in VTA.

**SIGNIFICANCE STATEMENT:** Opioid addiction is a chronically relapsing disorder defined by pathological drug-seeking behavior. Persistent relapse vulnerability is hypothesized to reflect a functional “rewiring” of reward-processing circuitry in the brain. This phenomenon is precipitated by drug-induced alterations in transcriptional plasticity of midbrain dopamine neurons, indicating an important role for epigenetic regulatory mechanisms in opioid dependence. We found that the newly discovered histone modification, H3 dopaminylation at glutamine 5 (H3Q5dop), was increased in ventral tegmental area (VTA) during periods of abstinence. H3Q5dop accumulation was found to regulate transcriptional programs associated with chronic heroin exposure. Using viral-mediated gene therapy to reduce H3Q5dop accumulation during abstinence resulted in attenuated heroin-primed seeking behaviors. Together, our results indicate that H3Q5dop plays an abstinence-specific role in regulating heroin-induced behavioral plasticity during prolonged abstinence.

## INTRODUCTION

Opioid Use Disorder (OUD) has dramatically increased to epidemic proportions in recent years, with heroin dependence growing more than 50% over the last decade and accounting for more drug-related deaths than any other substance of abuse [1-4]. Clinically, OUD is characterized by compulsive drug-seeking behaviors that can persist for years and extend over the course of a lifetime, with one of the highest risks of relapse among substance use disorders (SUD) [4]. Such vulnerability to relapse is a key diagnostic feature of addiction, and represents a highly complex process involving multiple circuits mediated by dynamic alterations in cellular plasticity within the mesolimbic reward system, where aberrant dopamine neurotransmission promotes maladaptive reward-processing and persistent deficits in motivated behaviors. Given the recurring, long-lasting nature of OUD, the field has begun to place great emphasis on drug-induced epigenetic regulation as a potential key mechanism in opioid addiction and relapse [5-9]. To date, however, our understanding of the causal mechanisms that mediate drug-induced transcriptional and cellular plasticity remains incomplete.

Recently, our laboratory demonstrated that in addition to modulating downstream synaptic activity as a neurotransmitter, dopamine itself plays a direct role in epigenetically coordinating gene expression within dopaminergic cells in ventral tegmental area (VTA) [10]. We previously found that certain monoamine neurotransmitters, including dopamine, can serve as donors for covalent modifications on specific residues of substrate proteins (termed monoaminylations) through an enzymatic process mediated by the writer-protein Transglutaminase 2 (TGM2) [11]. Among the identified substrate proteins for monoaminylation is histone H3, which is modified at glutamine 5 (Q5) on its N-terminal tail. At this H3Q5 site, specific histone monoaminylation marks (e.g., serotonylation or dopaminylation) were found to alter gene expression, either directly or through combinatorial interactions with other nearby histone PTMs, suggesting that these modifications may constitute an important epigenetic regulatory mechanism in monoaminergic neuromodulation [11, 12]. In a recent report, we confirmed that histone H3 dopaminylation (H3Q5dop) has a key functional role in cocaine-mediated transcriptional plasticity, dopaminergic neurotransmission, and cocaine-seeking behavior in a rodent model of drug relapse vulnerability [10]. Given the importance of H3Q5dop in mediating cocaine-induced transcriptional and behavioral plasticity, here we investigated whether functional roles for H3Q5dop represent a conserved mechanism across other addictive substances, or if this phenomenon was specific to cocaine.

All drugs of abuse are thought to elicit changes in dopaminergic neurotransmission from VTA, although different classes of substances act on this system through distinct cellular mechanisms. In addition, the magnitude and temporal dynamics of dopamine transmission vary widely between drugs, even within class, highlighting the importance of delineating substance-specific mechanisms in order to develop targeted therapeutic strategies to address the molecular changes associated with each drug [13]. In the present study, we investigated specific roles for H3Q5dop in heroin-induced transcriptional plasticity and drug-seeking behavior in a rodent model of relapse vulnerability. We found that prolonged abstinence from heroin SA resulted in the accumulation of H3Q5dop in VTA, though this increase occurs more rapidly in response to heroin *vs*. cocaine abstinence. We further showed that aberrant H3Q5dop dynamics in VTA contribute significantly to heroin-induced gene expression and drug-seeking behaviors. These findings indicate that H3Q5dop represents a conserved epigenetic substrate underlying addiction to distinct classes of abused drugs, though the gene expression programs that lead to altered DA output from VTA may be specific to heroin’s mechanism of action.

## RESULTS

### H3Q5dop is increased during both acute and prolonged abstinence in VTA following heroin SA

Given that histone dopaminylation (H3Q5dop, but not combinatorial H3 lysine 4 (K4) trimethyl (me3) Q5dop, was previously found to accumulate in VTA during prolonged, but not acute, abstinence from cocaine SA [10], we hypothesized that chronic exposure to heroin may similarly alter the deposition of H3Q5dop in this brain region. To assess this, we employed a heroin SA regimen wherein rats receive i.v. infusions of heroin or saline (as a non-reinforcement control) paired with a light cue (Fig. 1*A*) [14]. Following SA, rats underwent forced abstinence in their home cages for 1 d *vs*. 14 d, the latter of which represents a time period where cue-induced heroin seeking is expressed [15]. On abstinence day (AD) 1 or AD14, VTA tissues were collected and western blotting was performed to quantify histone dopaminylation levels (both H3Q5dop and H3K4me3Q5dop), as well as expression of an associated histone PTM (e.g., H3K4me3) and the writer Transglutaminase 2 enzyme. We found that H3Q5dop levels significantly increased (Fig. 1*B*) in VTA during both acute and prolonged abstinence from heroin SA. Neither H3K4me3Q5dop nor total H3K4me3 (Fig. 1*C*-*D*) showed increases at either time point. The writer enzyme for H3Q5dop was unaffected in its expression during periods of abstinence from heroin (Fig. 1*E*). Together, these data indicate that accumulation of histone dopaminylation in VTA occurs during long-term abstinence from heroin. Interestingly, heroin-induced accumulation of H3Q5dop was found to occur more rapidly after opioid exposure *vs*. cocaine, with significant increases detected as soon as AD1. This increase persisted to AD14, suggesting that H3Q5dop may contribute to opioid-induced transcriptional and behavioral alterations contributing to drug relapse.

**Figure 1.**
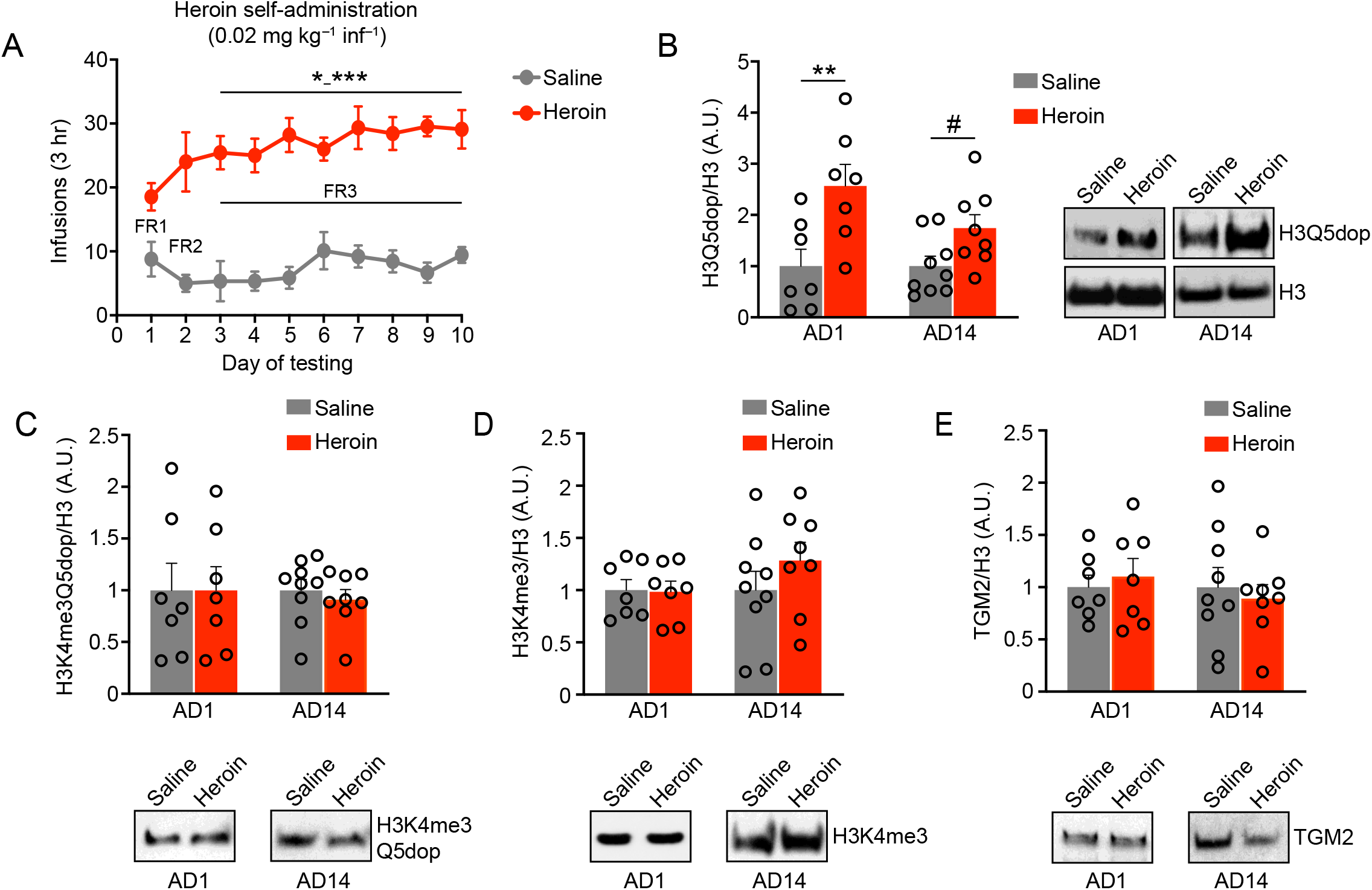
Histone H3 dopaminylation in VTA is dysregulated by heroin SA. (**A**) Number of infusions earned in daily 3 h (FR1 to FR3) test sessions in rats self-administering 0.02 mg/kg/infusion heroin or saline (*n*=8/group) for 10 days (two-way RM ANOVA, with Dunnett’s *post hoc* analysis *vs*. day 1, *p≤0.05 – ***p≤0.001). Western blot analysis of (**B**) H3Q5dop, (**C**) H3K4me3Q5dop, (**D**) H3K4me3 and (**E**) TGM2 in VTA tissues [abstinence day (AD) 1 *vs*. 14] from heroin *vs*. saline SA rats (*n*=7-9/group; two-way ANOVA with Sidak’s *post hoc* analysis, **p≤0.01, or an *a posteriori* Student’s *t* test, ^#^p≤0.05). All blots were normalized to total H3 as a loading control. Data presented as average ± SEM.

### H3Q5dop regulates heroin associated gene expression programs during prolonged abstinence

To explore the functional consequences of H3Q5dop accumulation during heroin abstinence, we used a lentiviral vector that expresses the H3 variant, H3.3, with a glutamine-to-alanine substitution at position 5 (H3.3Q5A), which cannot be dopaminylated. This H3.3Q5A vector has previously been shown to strongly express the mutant H3.3Q5A protein in a nuclear-specific manner in adult neurons, resulting in downregulation of H3Q5dop [10]. We delivered this H3.3QA vector into rodent VTA *vs*. H3.3 wildtype (WT) vs. empty vector controls [10, 11] in order to evaluate the impact of reducing H3Q5dop accumulation on gene expression programs during prolonged abstinence. After 10 d of heroin SA, rats were infected intra-VTA with one of the three viruses (on day 11), followed by a 30 d period of enforced abstinence to ensure maximal expression of viral transgenes (Fig. 2*A*), as previously described [10]. Following 30 d of forced abstinence, infected VTA tissues were microdissected and processed for RNA-seq. We assessed differential gene expression between H3Q5A *vs*. empty and between H3Q5A *vs*. WT viral groups. In both cases, attenuating H3Q5dop resulted in significant alterations in gene expression following drug SA.

**Figure 2.**
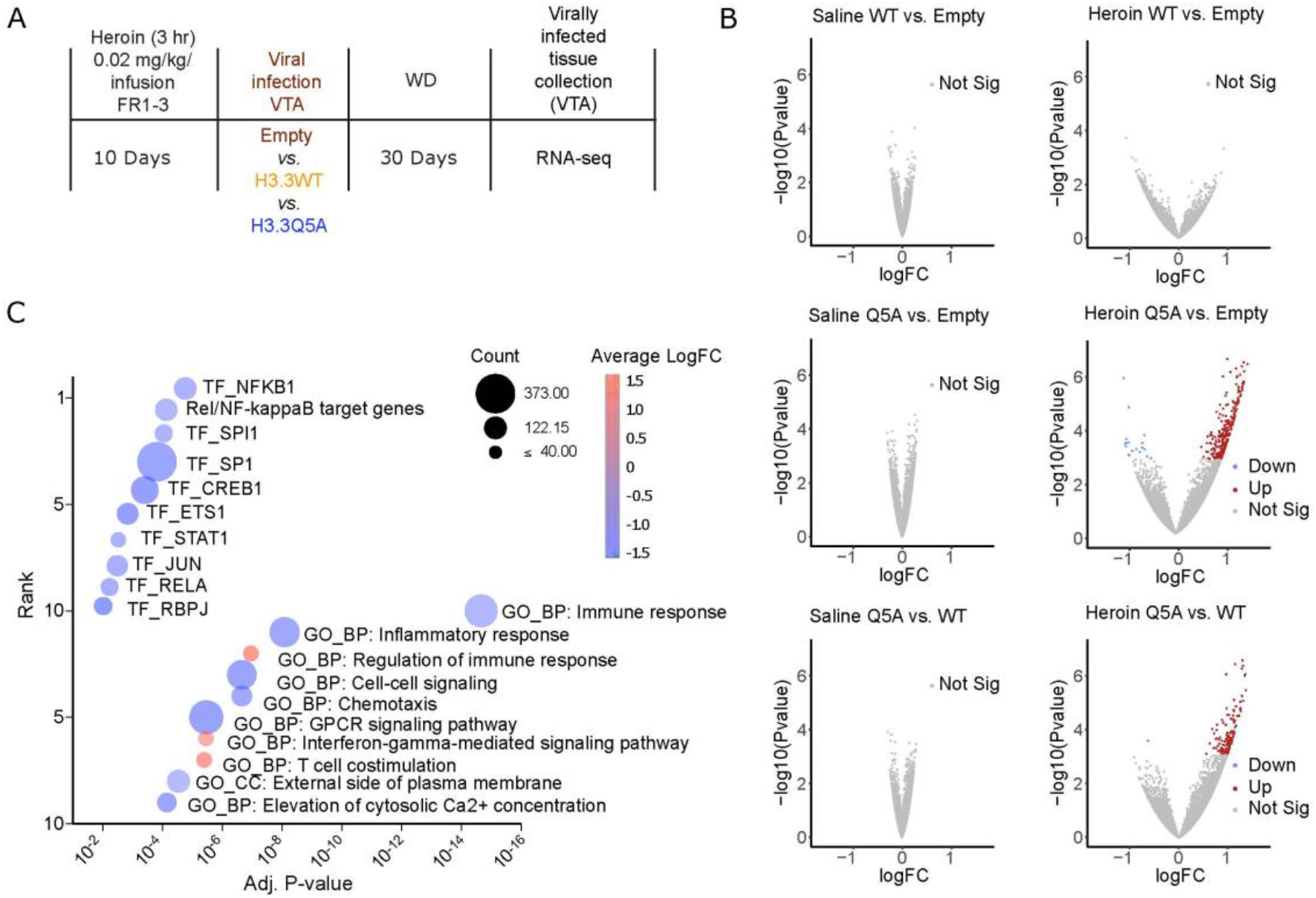
H3Q5dop in VTA contributes to heroin-mediated gene expression. (**A**) Experimental timeline of SA (heroin) RNA-seq experiment following viral transduction with either empty vector or H3.3 WT *vs*. H3.3Q5A viruses (*n*=4-7/group). (**B**) Volcano scatter plots display −log10(p-value) *vs*. logFC gene expression for each pairwise comparison shown: H3.3Q5A *vs*. empty or H3.3WT, and H3.3WT *vs*. empty groups following either heroin SA or saline SA (the latter data was extracted from [10]). Color points indicate FDR<0.1 (**C**) Gene Set Enrichment Analyses for the pairwise differential expression comparison between H3.3Q5A *vs*. H3.3WT. Multivariant bubble plots display the top 10 gene sets identified, Y-axis = rank from 1-10 for Hallmark and GO gene sets from GeneSetDB based upon significance score, plotted according to adj. p-value (X-axis), the direction of regulation (fill color), and the number of DEx genes in the gene set.

We first employed pairwise comparisons to identify differentially expressed (DEx) genes between H3.3Q5A and control viral vectors post-heroin SA. We identified 268 and 131 DEx genes in the H3.3Q5A *vs*. empty or H3.3 WT comparisons, respectively, at a false discovery rate (FDR) of < 0.1; Fig. 2*B* and *SI Appendix*, Tables S1-2). These genes were primarily upregulated in their expression (Fig. 2*C*). Note that 110 of these genes were DEx in both comparisons, and furthermore, there were no DEx genes between empty *vs*. H3.3, confirming that the majority of gene expression changes induced by the H3Q5A mutant are not due to exogenous expression of the histone H3.3 protein. In order to determine whether these H3.3Q5A mediated alterations were heroin-specific events, we also examined the gene expression profile of H3.3Q5A *vs*. empty or H3.3WT in the context of saline SA, where we found no DEx genes regulated by the dominant negative virus (Fig. 2*B*), indicating these changes are dependent on prior heroin exposures. We next performed Gene Set Enrichment Analysis (GSEA) on the normalized expression matrix comparing transcript counts between the H3.3Q5A and H3.3 WT conditions and found significant gene set associations [Gene Regulation (GR) and Gene Ontology (GO)] with biological processes involved in the regulation of immune signaling (e.g., Immune Response, Inflammatory Response), cellular communication (GPCR pathways, cell signaling, CA2+ elevation), and transcriptional regulation of stress response and transcription regulation (e.g., NF-kB, RelA, and Jun) (Fig. 2*C)*. Interestingly, the genes within these inflammatory sets were downregulated by Q5A, indicating that H3Q5dop accumulation during heroin abstinence may work to potentiate immune activation stress.

These findings suggest that H3Q5dop plays an important role in heroin-induced transcriptional plasticity in VTA, similar to that observed with cocaine [10]. However, the pathways regulated by H3 dopaminylation in the context of heroin were distinct from those mediated by cocaine, which primarily overlapped with gene set categories associated with synaptic transmission and long-term potentiation, indicating a more direct effect on neuronal signaling. Heroin-induced gene expression was significantly associated with gene sets linked to regulation of inflammatory response, confirming previous reports that this drug has critical immunomodulatory consequences.

### Accumulation of H3Q5dop in VTA contributes to heroin-seeking in a drug-primed reinstatement paradigm

Finally, to investigate the consequences of reducing H3Q5dop levels on heroin relapse-relevant behaviors, we put an independent cohort of animals through a chronic heroin self-administration paradigm, followed by intra-VTA viral infusion on AD1, using the same three viral vectors discussed above (Fig. 3*A*-*B*). On AD30, rats were extinguished to the drug-paired context over two daily sessions and were subsequently tested for drug-primed heroin-seeking behavior. Preventing H3Q5dop accumulation in heroin SA animals significantly reduced drug-induced reinstatement of motivated behaviors (Fig. 3*C*) without altering locomotor functions (*SI Appendix*, Fig. S1*A*). This indicates that aberrant accumulation of H3 dopaminylation in VTA represents a discrete transcriptional program mediating relapse associated vulnerability across pharmacologically distinct addictive substances.

**Figure 3.**
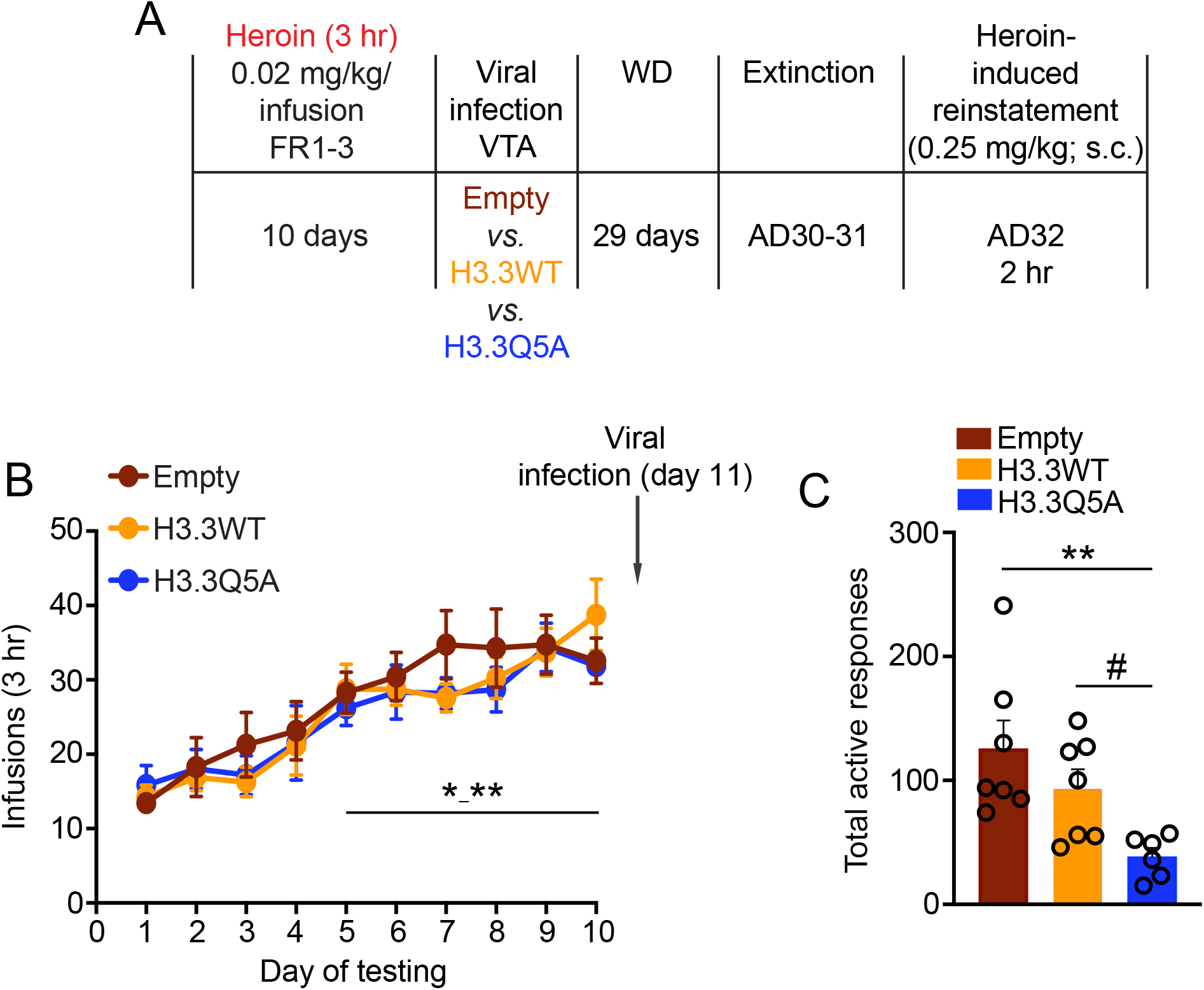
Reducing H3Q5dop in VTA attenuates heroin-seeking. (**A**) Experimental timeline of heroin SA drug-induced reinstatement experiments following viral transduction with either empty vector, H3.3 WT or H3.3Q5A viruses. (**B**) After 10 days of heroin SA [with escalation of intake observed, *n*=10/group, empty vector and H3.3 WT *vs*. H3.3Q5A (heroin): two-way RM ANOVA, with Dunnett’s *post hoc* analysis *vs*. day 1, *p≤0.05 – **p≤0.01], rats were infected intra-VTA with one of the three viral vectors (on day 11), followed by 30 d of withdrawal, extinction and (**C**) drug-induced reinstatement (*n*=6-7/group); the total number of active responses was reduced in H3.3Q5A rats *vs*. H3.3 WT and empty vector controls (one-way ANOVA with Tukey’s *post hoc* analysis, **p≤0.01 or an *a posteriori* Student’s *t* test, ^#^p≤0.05). Data presented as average ± SEM.

## DISCUSSION

As part of its mechanism of action, heroin exposures increase neurotransmission from dopaminergic neurons in VTA to reward-processing projection regions in striatum. These fluctuations in dopamine signaling are thought to contribute to maladaptive plasticity and impaired decision making observed during substance dependence. However, the mechanisms that mediate these processes at the epigenetic level are not fully characterized. Here, we showed that H3Q5dop rapidly accumulates in rodent VTA at 24 hours post-heroin SA, and this increase persisted during protracted periods of heroin abstinence. We next profiled gene expression changes that occur after attenuating levels of H3Q5dop during heroin abstinence and identified significant overlap with regulation of inflammatory signaling and NF-kB pathways, with H3Q5A animals showing reduction of genes in these categories. Furthermore, these expression changes coincided with a significant decrease in drug-seeking behavior after 30 d of heroin abstinence. Together, these data indicate that H3Q5dop plays a critical role in promoting persistent functional alterations that drive maladaptive behaviors related to heroin relapse vulnerability.

Our findings highlight the convergent role of H3Q5dop in regulating epigenetic changes after chronic exposures and abstinence from different drugs of abuse (e.g., opioids *vs*. psychostimulants). However, we also observed key differences between the substances, with respect to both the dynamics and gene targets of H3Q5dop. We found that H3Q5dop is significantly increased in rodent VTA as soon as AD1 post-heroin SA, an increase that persists throughout chronic abstinence. In contrast, levels of H3Q5dop during cocaine abstinence remained unchanged 24 hr after the last day of cocaine SA, and then slowly accumulated during long-term withdrawal. This temporal difference in H3Q5dop dynamics may reflect differences in dopaminergic activity in response to either substance, particularly during more acute periods of abstinence – for example, electrophysiology and fiber photometry studies have demonstrated that while cocaine exposures lead to immediate inhibition of dopaminergic VTA neurons, heroin exposures strongly activate these projections [16]. Furthermore, neurons in striatum activate distinct pathways in downstream projection regions in response to heroin *vs*. cocaine, suggesting that the reward system may display circuit-specific responses to different classes of drugs of abuse [17, 18].

The second major difference revealed in our current study relates to the VTA gene expression changes mediated by H3Q5dop in response to heroin *vs*. cocaine SA. Gene set enrichment analyses revealed a significant enrichment of gene sets associated with inflammatory responses and activation of NF-kB processes – however in the H3Q5A conditions, genes in these sets were downregulated compared to empty or H3.3WT groups. These findings are particularly relevant to the well-characterized immunomodulatory effects of chronic heroin use, which have been shown to play a central role in heroin induced neuroplasticity [19, 20]. Mounting evidence has revealed that opioid dependence suppresses adaptive immune signaling, increases pro-inflammatory NF-kB pathways, and activates both central and peripheral mechanisms of reactive stress responses, with expression levels of these factors correlating positively with drug-taking in animal models of heroin use [21-23]. Given that chronic heroin use increases feed-forward inflammatory NF-kB signaling [24, 25], it is notable that reducing H3Q5dop acts to specifically downregulate this pathway, as well as other innate immune responses, suggesting that this modification may actively participate in the molecular translation of heroin withdrawal to altered inflammatory transcription.

Finally, our previous work demonstrated that the H3Q5dop modification may be an upstream epigenetic regulator of dopaminergic neurotransmission in drug dependence. Our current findings indicate that manipulation of H3Q5dop during different stages of drug dependence may represent a useful tool to reveal important gene expression profiles that are unique to specific classes of substances. Moving forward, delineating specific transcriptional signatures that effectively promote reward-circuit remodeling for different drugs of abuse may provide insights to guide clinicians in tailoring therapeutic strategies towards the unique mechanisms of action for a given drug class. For example, substances like nicotine and alcohol, which also converge on promoting dopamine outflow, have distinct molecular interactions with cellular substrates in the VTA. Further study of H3Q5dop dynamics in the context of these different substances may reveal distinct epigenetic mechanisms that translate their respective receptor-binding signals into increased plasticity. Overall, the data presented here demonstrate that different classes of abused substances act through H3Q5dop deposition as a convergent epigenetic mechanism to recruit specific transcriptional programs that alter dopaminergic neurotransmission and drive drug-seeking behaviors during prolonged abstinence.

## MATERIALS AND METHODS

### Animals

Male Sprague-Dawley rats (Envigo; 250-300 g) were pair housed and maintained on a 12 h reverse light/dark cycle with food and water available *ad libitum*. All procedures were done in accordance with NIH guidelines and the Institutional Animal Care and Use Committees of the Icahn School of Medicine at Mount Sinai and the State University of New York at Buffalo.

### Jugular Catheterization

For heroin SA, rats were implanted with chronic indwelling jugular catheters, as previously reported [14, 26]. Rats were given 5 d of recovery, during which time the catheters were flushed with 0.2 mL of a solution containing enrofloxacin (4 mg/mL) and heparin saline (50 IU/mL in 0.9% sterile saline) to preserve catheter patency. All animals received an i.v. infusion of Ketamine hydrochloride (0.5 mg/kg in 0.05 mL) one day prior to behavioral training to verify catheter patency. Rats that responded with a loss of muscle tone and righting reflexes were considered patent and were included in cohorts for behavioral training.

### Heroin Self-Administration and Viral surgeries

Rats were allowed to recover from jugular catheterization surgery for 5 d, and then were trained to SA heroin (0.02 mg/kg/inf) or saline in operant chambers for 3 h each day for a period of 10 d. A response in the active hole nose poke resulted in an intravenous injection of heroin (0.02 mg/kg/inf) or saline using a fixed ratio 1 (FR1) schedule of reinforcement on day 1 that was increased to FR3 on day 3 and maintained for the remainder of the experiment [14]. Responses in the inactive hole resulted in no programmed consequences. Following training sessions, catheters were flushed, and rats were returned to their colony rooms. For viral manipulations, all rats self-administered heroin and were bilaterally injected intra-VTA with either lenti-H3.3WT or lenti-H3.3Q5A (*vs*. an empty vector control) on AD11 (see below for additional information on surgery coordinates).

### Drug-induced Reinstatement

For drug-induced reinstatement, on AD30, rats were returned to experimental chambers for a within-session extinction procedure, as described elsewhere [14]. Briefly, rats were allowed to respond for 8-10 1 h sessions in the presence of house light and cues with 5-min intervals, during which times the house light was extinguished. At 24 h following the extinction session, when responses fell to less than 30 per session, rats were injected with 0.25 mg kg−1 of heroin (s.c.) and returned to the operant chambers for 2 h to respond for drug-induced reinstatement.

### Western Blotting and Antibodies

VTA tissues were collected from heroin SA rats (2 mm punches) on AD1 or AD14 and frozen. Nuclear fractions were purified via homogenization in Buffer A containing 10 mM HEPES (pH 7.9), 10 mM KCl, 1.5 mM MgCl_2_, 0.34 M sucrose, 10% glycerol, 1 mM EDTA, 1X protease inhibitor cocktail and 1X phosphostop inhibitor. After homogenization, 0.1% TritonX-100 was added, samples were incubated on ice for 30 min and then centrifuged for 5 min at 1300g at 4° C. Supernatants (i.e., cytosolic fractions) were discarded, and nuclear pellets were resuspended in Buffer A to remove any remaining cytosolic contamination, followed by centrifugation for 5 min at 1300g at 4° C. Supernatants were then discarded and pellets resuspended and sonicated in sample buffer containing 0.3 M sucrose, 5 mM HEPES, 1% SDS and 1X protease inhibitor cocktail. Protein concentrations were measured with the DC protein assay kit (BioRad), and 10-15 ug of protein was loaded onto 4-12% NuPage BisTris gels (Invitrogen) for electrophoresis. Proteins were next transferred to nitrocellulose membranes and blocked for 1 h in 5% milk in PBS + 0.1% Tween 20 (PBST), followed by primary antibody incubation for two days at 4° C. The following antibodies were used in this manuscript: rabbit anti-H3Q5dop (1:200, ABE2588; Millipore), rabbit anti-H3K4me3Q5dop (1:500, ABE2590; Millipore), rabbit anti-H3K4me3 (1:1000, lot #: GR273043-6, Abcam ab8580), mouse anti-Tgm2 (1:500, lot #: Gr3188112-4, Abcam ab2386) and rabbit anti-H3 (1:50000, lot #: GR293151-1, Abcam ab1791). Membranes were then washed 3X in PBST (10 min) and incubated for 1 h with horseradish peroxidase conjugated anti-rabbit (BioRad 170-6515, lot #: 64033820) or anti-mouse (GE Healthcare UK Limited NA931V, lot #: 9814763) secondary antibodies (1:10000; 1:50000 for anti-H3 antibody, BioRad) in 5% milk/PBST at RT. After three more washes with PBST, bands were detected using enhanced chemiluminescence (ECL; Millipore). Densitometry was used to quantify protein bands using Image J Software (NIH), and proteins were normalized to total H3.

### Lentiviral Constructs and Viral Infusion

Lentiviral H3.3 constructs [empty *vs*. wildtype (WT) *vs*. (Q5A)-Flag-HA] were generated and validated as previously described [10, 11].The H3.3Q5A vector used in these studies, which does not affect adjacent H3K4me3, inhibits the expression of all modifications on Q5. However, appreciable signal for other monoaminyl marks (e.g., H3Q5ser) is not observed in VTA *vs*. H3Q5dop [10]. After 24 hr following the last day of heroin *vs*. saline SA (day 11), animals were anaesthetized with a ketamine/xylazine solution (80/6 mg/kg) i.p., positioned in a stereotaxic frame (Kopf instruments) and 1 μl of viral construct was infused bilaterally into VTA using the following coordinates; anterior-posterior (AP) −4.9, medial-lateral (ML) +2.1, dorsal-ventral (DV) −7.6. After surgery, rats received meloxicam (1 mg/kg) s.c. and topical antibiotic treatments for ∼3 days. All tissue collections or behavioral testing commenced 30 d after viral surgery to allow for maximal expression of the viral constructs, as previously validated [10].

### RNA Isolation and Sequencing

Rat brains were removed and immediately snap frozen. Brains were then sectioned into 100 uM thick coronal slices on a cryostat at −20° C, and 2 mm punches were dissected from virally infected VTA using RFP-fluorescence illuminated via a NIGHTSEA Dual Fluorescent Protein flashlight (#DFP-1). Total RNA was then extracted from punches using Trizol (Thermo Fisher #15596026)-chloroform, followed by cleanup and elution into RNAase-free water using Qiagen minelute (#74204) columns, as per manufacturer instructions. Following elution, RNA samples were enriched for mRNA via polyA tail selection beads, and mRNA libraries were prepared using the Illumina Truseq RNA Library Prep Kit V2 (#RS-122-2001). Libraries were pooled and sequenced on the Illumina Novaseq platform. RNA-seq data were pre-processed and analyzed, as previously described [10]. Briefly, FastQC (Version 0.72) was performed on the concatenated replicate raw sequencing reads from each library to ensure minimal PCR duplication and sequencing quality. Reads were aligned to the rn6 genome using HISAT2 (Version 2.1.0) and annotated against Ensembl v90. Multiple-aligned reads were removed, and remaining transcript reads were counted using featurecounts (Version 2.0.1). DESEQ2 [27] (Version 2.11.40.6) was then used to normalize read counts between the three viral groups, and to perform pairwise differential expression analyses. Differentially expressed (DEx) genes were defined at FDR cutoff < 0.1. DEx genes between Heroin H3.3Q5A vs. RFP empty vector (268) were overlapped (Venny) with DEx genes between Heroin H3.3Q5A vs. H3.3 WT groups (131) to identify the subset of shared DEx genes between these comparisons (110).

In order to visualize genes that display large magnitude fold changes as well as statistical significance (FDR <0.1), a volcano scatter plot was constructed for each pairwise comparison shown, using a galaxy implementation of ggplot2 (Version 0.0.3). The points were plotted according to −log10(p-value) on the Y-axis, and logFC on the X-axis, derived from the DESEQ2 output. Processed read count data for Saline comparisons was extracted from our previous publication [10], deposited in the National Center for Biotechnology Information Gene Expression Omnibus database under accession number GSE124055. In order to determine if differential gene expression comparisons showed any significant overlaps with annotated gene sets, Gene Set Enrichment Analyses was performed using the EGSEA package (Version 1.10.0) against EGSEA’s gene set database (KEGG and GeneSetDB). Results from six algorithms (camera, safe, gage, gsva, globaltest, ora) were used to calculate collective gene set scores, and gene sets were ranked by adj. p-value [28]. Multivariant plots display top 10 gene sets, Y-axis = rank from 1-10 for Gene Regulation (GR) and Gene ontology (GO) gene sets from GeneSetDB based on significance score, plotted according to p-value (X-axis), number of DEx genes overlapping with the gene set (size), and the direction of regulation (fill color).

### Statistical Analyses

For all behavioral testing and molecular experiments involving more than two treatments and time points, two-way or one-way ANOVAs were performed with subsequent Sidak’s, Tukey’s or Dunnett’s *post hoc* analyses, and/or *a posteriori* t*-*tests (as indicated throughout the text). In molecular analyses, all animals used were included as separate *n*s (samples were not pooled). Significance was determined at p<0.05. All data are represented as mean ± SEM.

## ACKNOWLEDGEMENTS

The NIDA Drug Supply Program generously gifted the heroin used in these studies. This work was supported by grants from the National Institutes of Health: DP1 DA042078 (I.M.), R01 DA015521 (D.M.D.), R21 DA0488554 (D.M.D.), U01 DA051947 (D.M.D.), F31 MH124425 (S.L.F.), F31 DA045428 (A.E.L) and F99 NS108543 (J.A.M.), as well as awards from: Brain Research Foundation’s Fay/Frank Seed Grant Award (I.M.).

***SI Appendix***

**Figure S1.**
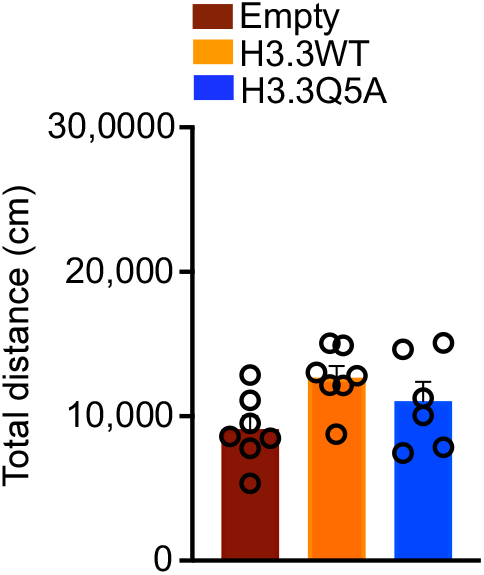
Reducing H3Q5dop in VTA does not impact general locomotion. Total locomotion during drug-induced reinstatement (*n*=6-7/group); the total distance moved (cm) was not affected in H3.3Q5A rats *vs*. H3.3 WT or empty vector controls (one-way ANOVA, p>0.05. Data presented as average ± SEM.

Supplementary Data **Tables S1-2** (Processed RNA-seq Data; Excel Format)

